# Early signals of vaccine driven perturbation seen in pneumococcal carriage population genomic data

**DOI:** 10.1101/459693

**Authors:** Chrispin Chaguza, Ellen Heinsbroek, Rebecca A. Gladstone, Terence Tafatatha, Maaike Alaerts, Chikondi Peno, Jennifer E. Cornick, Patrick Musicha, Naor Bar-Zeev, Arox Kamng’ona, Aras Kadioglu, Lesley Mcgee, William P. Hanage, Robert F. Breiman, Robert S. Heyderman, Neil French, Dean B. Everett, Stephen D. Bentley

**Affiliations:** Parasites and Microbes, Wellcome Sanger Institute, Wellcome Genome Campus, Cambridge, UK; Department of Clinical Infection, Microbiology and Immunology, Institute of Infection and Global Health, University of Liverpool, Liverpool, UK; Malawi-Liverpool-Wellcome Trust Clinical Research Programme, Blantyre, Malawi; HIV & STI Department, National Infection Service, Public Health England, London, UK; Malawi Epidemiology Intervention Research Unit (formerly KPS), Chilumba, Malawi; Center of Medical Genetics, University of Antwerp, Antwerp, Belgium; MRC Centre for Inflammation Research, Queens Medical Research Institute, University of Edinburgh, Edinburgh, UK; Mahidol Oxford Tropical Medicine Research Unit, Mahidol University, Bangkok, Thailand; Nuffield Department of Medicine, University of Oxford, Oxford, UK; Department of International Health, Johns Hopkins Bloomberg School of Public Health, Baltimore, USA; Department of Biomedical Sciences, University of Malawi, College of Medicine, Blantyre, Malawi; Respiratory Diseases Branch, Centers for Disease Control and Prevention, Atlanta, USA; Center for Communicable Disease Dynamics, Department of Epidemiology, Harvard T.H. Chan School of Public Health, Boston, Massachusetts, USA; Hubert Department of Global Health, Rollins School of Public Health, Emory University, Atlanta, USA; Division of Infection and Immunity, University College London, London, UK; Department of Pathology, University of Cambridge, Cambridge, UK

## Abstract

Pneumococcal conjugate vaccines (PCV) have reduced pneumococcal diseases globally. Despite this, much remains to be learned about their effect on pathogen population structure. Here we undertook whole genome sequencing of 660 pneumococcal strains from asymptomatic carriers to investigate population restructuring in pneumococcal strains sampled before and after PCV13 introduction in a previously vaccine-naïve setting. We show substantial decreasing frequency of vaccine-type (VT) strains and their strain diversity post-vaccination in the vaccinated but not unvaccinated age groups indicative of direct but limited or delayed indirect effect of vaccination. Clearance of identical VT serotypes associated with multiple lineages occurred regardless of their genetic background. Interestingly, despite the increasing frequency of non-vaccine type (NVT) strains through serotype replacement, the serotype diversity was not fully restored to the levels observed prior to vaccination implying limited serotype replacement. The frequency of antibiotic resistant strains was low and remained largely unchanged post-vaccination but intermediate-penicillin-resistant lineages were reduced in the post vaccine population. Significant perturbations marked by changing frequency of accessory genes associated with diverse functions especially mobile genetic elements and bacteriocin activity were detected. This phylogenomic analysis demonstrates early vaccine-induced pneumococcal population restructuring not only at serotype but also accessory genome level.

**Author summary:** Different formulations of PCVs have been effective in reducing the invasive pneumococcal disease burden globally. Clinical trials have started to indicate high impact and effectiveness of PCV13 in Sub Saharan Africa (SSA) but there is limited understanding of how the introduction of PCVs alters the population structure of pneumococcal strains at serotype and genomic level. Here we investigated this using pneumococcal strains sampled pre‐ and post-PCV13 introduction from a previously vaccine naïve setting in Northern Malawi. Our findings reveal decrease in frequency of VT serotypes and their associated lineages in the largely vaccinated under-five population but not older individuals indicating a direct but limited or delayed indirect protection. The diversity of serotypes also decreased post-vaccination in VT strains in the under-fives but there was no change in NVT strains suggesting incomplete serotype replacement. At the genomic level, logistic regression revealed changing frequency of accessory genes largely associated with mobile genetic elements but such changes did not include any antibiotic resistance genes. These findings show significant perturbations at serotype and accessory genome level in carried pneumococcal population after two years from PCV13 introduction but the pneumococcal population was still perturbed and had not returned to a new equilibrium state.

## Introduction

Pneumococcal polysaccharide antigens covalently attached to carrier proteins elicit sufficient serotype-specific antibody responses against *Streptococcus pneumoniae* (the pneumococcus) and form the basis of pneumococcal conjugate vaccines (PCV). Different formulations of PCVs have been licensed and introduced globally, and clinical case-control, cohort and surveillance studies have documented high effectiveness of these vaccines on non-invasive [1] and invasive pneumococcal disease (IPD) [2]. For example, after introduction of PCV7, 69% and 57% reductions in IPD were observed in USA and UK respectively whereby vaccine type (VT) strains decreased by >90% [2, 3]. Few African countries introduced PCV7 because its projected low coverage of VTs as highly common invasive serotypes in this setting (e.g. serotypes 1 and 5) were not covered [4]. Regardless, PCV7 caused >85% reduction in IPD in South Africa, crucially in HIV-infected individuals [5]. Consistent with findings in high-income countries [6], higher-valent PCVs appears to be highly effective in reducing VT serotypes in carriage (>65%) and IPD (>80%) in Africa[7–9].

In addition to reducing IPD [2, 8, 9], PCV has an added benefit of reducing VT carriage [10]. However, the impact on overall carriage rate and density is not substantial [11, 12]. This is because PCV introduction induces serotype replacement whereby reduction of VTs substantial alters serotype competition dynamics thereby prompting an increase of non-vaccine type (NVT) strains uncommon prior to vaccination [13]. A well-known example of replacement is the upsurge of serotype 19A post-PCV7 introduction [14]. While invasive potential of replacement NVT serotypes is typically lower than for VTs, this is not universally true, and some strains retain propensity for invasive disease [15]. Therefore, understanding post-vaccination dynamics in pneumococcal population is crucial to inform future clinical interventions and it may be more informative to study the carriage population where evolution is ongoing, rather than studying isolates from disease which is considered to be an evolutionary dead-end.

In this study, we undertook whole genome sequencing (WGS) of 660 pneumococcal strains sampled before and after nation-wide introduction of PCV13 vaccine in a previously vaccine-naïve setting of Northern Malawi to investigate early changes in population structure, serotype composition and diversity, accessory genome variation. PCV13 was introduced in November 2011 in Malawi was via an accelerated ‘3+0’ schedule (6, 10 and 14 weeks) with a limited catch-up for infants in the first year from introduction [16]. WGS of the isolates was undertaken by the Global Pneumococcal Sequencing (GPS) project (www.pneumogen.net), which to date has sequenced ≈23,000 pneumococcal strains sampled globally to map out post-vaccination evolution patterns of strains to inform future vaccine design. Here we focus on carriage samples from northern Malawi and present evidence of early vaccine-induced population restructuring at serotype, lineage and accessory genome level.

## Results

### Defining the population structure

Genomes were sequenced for 660 isolates collected from healthy carriers before and after introduction of PCV13 in a previously vaccine-naïve population in the Karonga district of northern Malawi (Fig 1a, S1 Data). A total of 45 serotypes and 169 sequence types (STs) were detected with evidence of serotype-switching within lineage due to recombination [17]. From a reference-free 1,050,021bp multiple sequence alignment of the 660 study genomes we identified 88,961 single nucleotide polymorphisms (SNP) which were used to infer their genetic population structure using an unsupervised hierarchical clustering algorithm [18] (S1-5 Fig). This approach identified twenty-three genomic clusters (GC) or lineages with one GC (GC23) appearing to be polyphyletic (Fig 1b). The GCs exhibited different within-lineage sequence diversity due to variations in recombination and age of the lineage (S6 Fig). We then used this defined population to investigate changes in frequency of serotypes, lineages and accessory genes post-PCV13 introduction while controlling for serotype category and sampling period.

**Fig 1:**
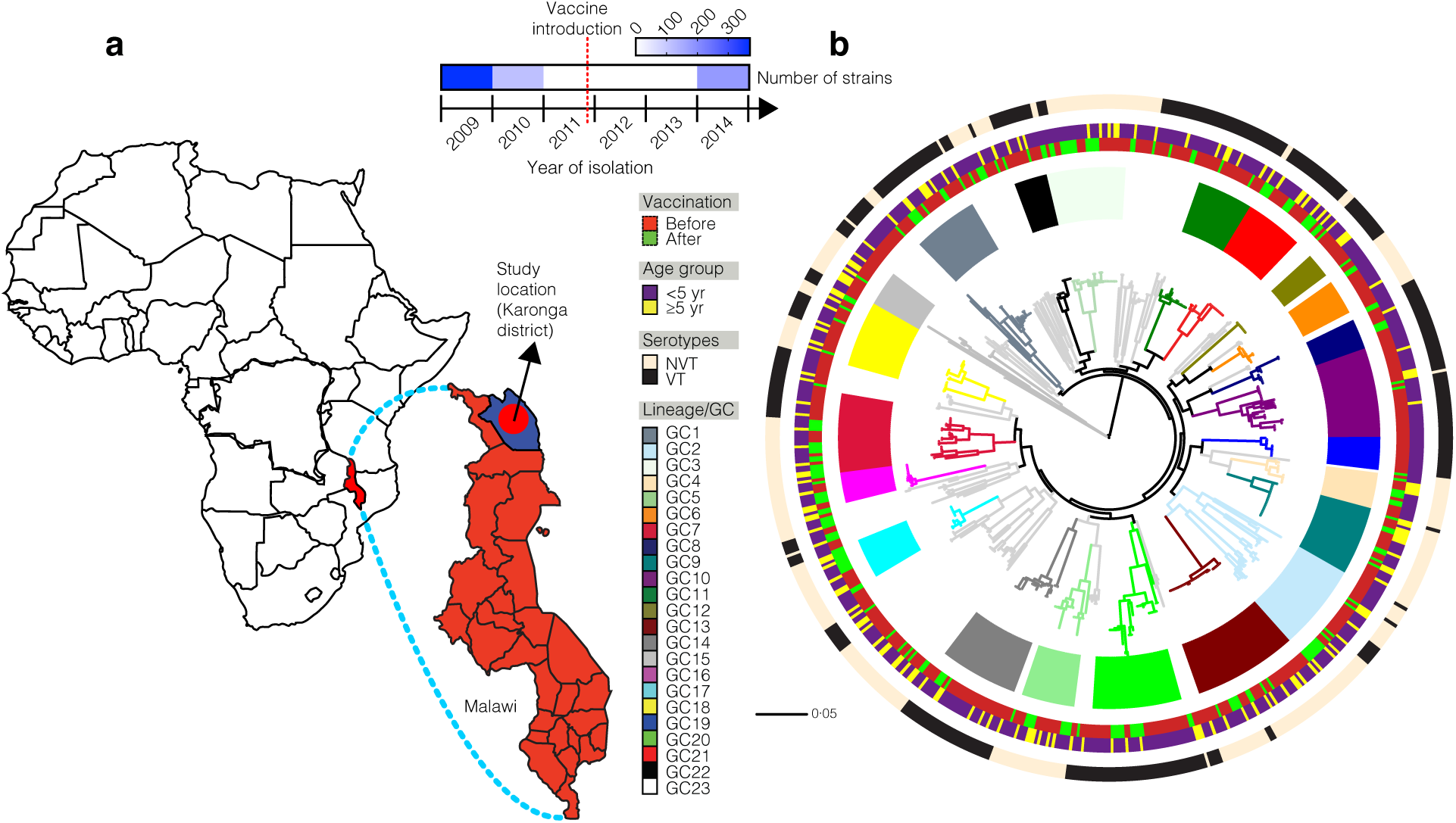
Core gene maximum likelihood phylogeny of carried pneumococci. (a) Sampling location of the study strains. (b) Maximum likelihood phylogeny of the carried pneumococcal strains reconstructed using core genome SNPs from 660 study strains demonstrating their genetic similarity and diversity. The strains’ metadata namely GC (lineage), sampling period, age group and serotype category of the subjects shown by coloured strips around the phylogeny. The colours for phylogenetic branches correspond to the inferred GCs as shown in the metadata key next to the tree. The tree was rooted at the mid-point of the branch separating ‘classical’ non-typeable (NT) pneumococcal GC (GC15) from rest of the strains. Detailed characteristics of the study strains are provided in S1 Data.

### Decrease of VT serotypes and their GCs signify PCV impact

Changes in frequency of GCs and their constituent serotypes before and after vaccination signaled substantial restructuring of the pneumococcal carriage population two years after PCV introduction. Using the Fisher’s Exact test, three GCs namely GC3 (*P*=0.002), GC16 (*P*=0.004) and GC17 (*P*=0.004) showed a post-vaccination increase in frequency but there was a decrease of GC10 (*P*=0.016) and GC19 (*P*=0.026) (Fig 2a,b, S1 Table). The reduced frequency of GCs, which were highly associated with VTs, largely reflected reduction of VTs in vaccinated children under five years old; a reduction that was not seen in the over five year old unvaccinated population (Fig 2c,d, S2 Table). We modelled the relationship between frequency of VT and NVT serotypes pre‐ and post-vaccination using linear regression and there was a smaller regression coefficient for VTs than NVT serotypes (Fig 2c,d). This showed that PCV13 reduced frequency of VT at a higher rate than NVT serotypes in the GCs. The majority of the GCs showed no change in odds ratio of VT serotypes in under-fives relative to over-fives while only four GCs (GC4,16,19,22) showed significant changes post-vaccination (S7 Fig, S3 Table). The decrease in the overall frequency of VT serotypes was evident (*P*=4.80 ×10^−8^, Fisher’s Exact test) in strains sampled from under-fives but not over-fives (*P*=0.3739, Fisher’s Exact test) (Fig 2e-g). The frequency of VT strains was also higher in strains from under-fives than over-fives (*P*=8.44×10^−4^, Fisher’s Exact test) but following vaccination their frequency became equal (33%) in both age groups *(*P*=1*, Fisher’s Exact test) (Fig 2g, S4,5 Table). Similar tests revealed significant increase of four NVTs serotype 7C (*P*=0.001), 15B/C (*P*=0.004), 23A (*P*=0.017) and 28F (*P*=0.0001) after vaccination in under-fives while only 28F (*P*=0.029) increased in over-fives. By using a unique collection of isolates from both children and adults pre‐ and post-vaccination, these findings demonstrate substantial direct effects of PCV in children but either limited or incomplete indirect protection against carriage of VT strains in older individuals.

**Fig 2:**
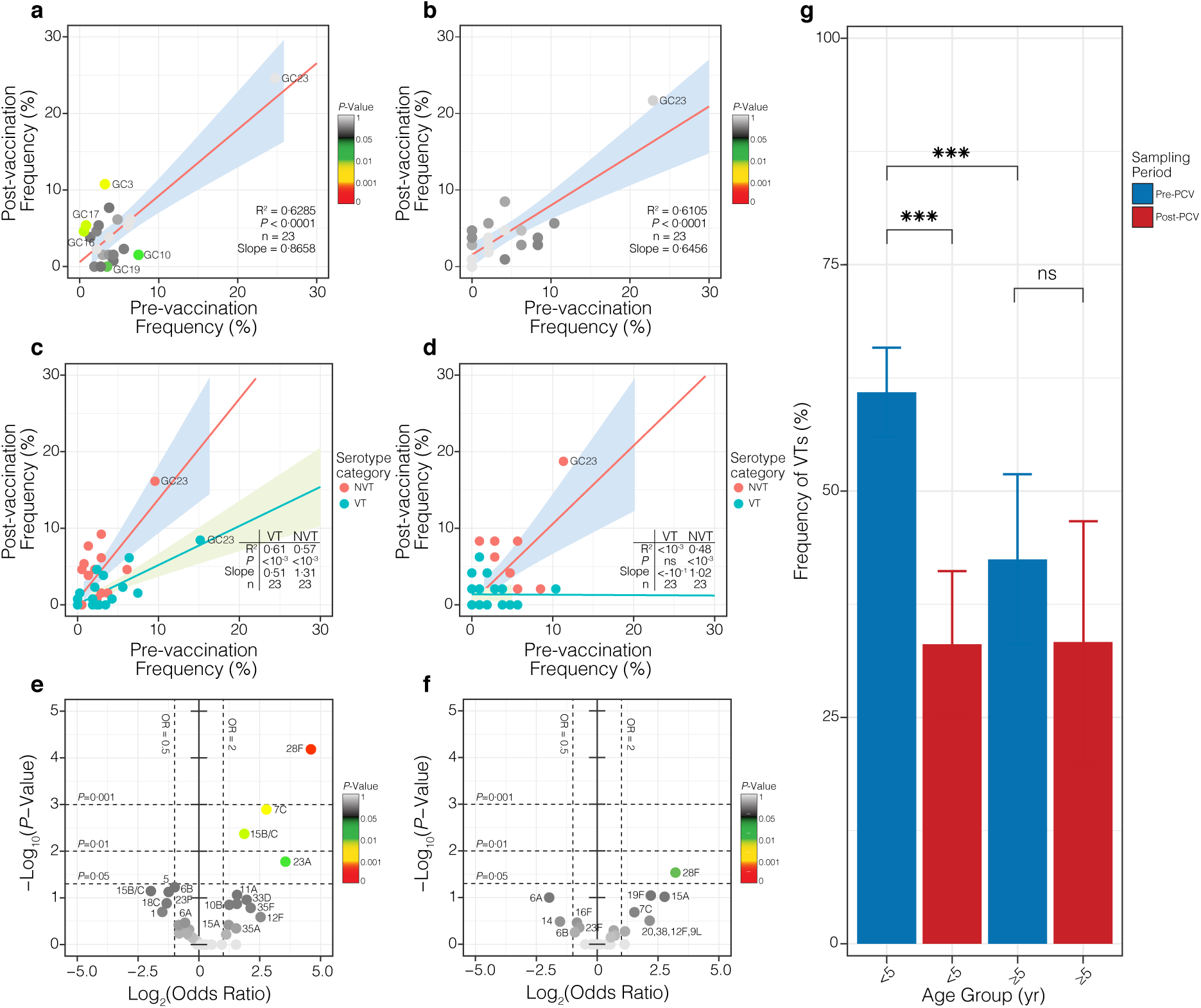
Frequency of GCs and serotypes in carriage. The scatter plot showing frequency of GCs before and after vaccination in (a) under-fives and (b) over-fives. Fitted linear regression for the overall trend in GC frequency of VT and NVT strains pre‐ and post-vaccination. (c) The scatter plots showing frequency of VT serotypes in GCs pre‐ and post-vaccination in (d) under-fives and (e) over-fives. The regression lines are bordered by 95% CI. The changes of individual serotype frequency are shown by volcano plots for strains in (f) under-fives and (g) over-fives where the x-axis shows magnitude (log_2_ odds ratio) and y-axis shows statistical significance (log_10_ *P*-value) of change after compared to before vaccination. Statistically significant changes are marked with asterisks: ‘ns’: not significant, *P*<0.05 (*), *P*<0.01 (**) and *P*<0.001 (***).

### Emergence and clonal expansion of NVT strains

The overall frequency of serotypes and their dynamics within GCs revealed clonal expansion and potential emergence through capsule switching (Fig 4). With the exception of serotype 28F, all serotypes were detected prior to vaccination suggesting that emergence of previously undetected serotypes following vaccination was uncommon. Therefore, clonal expansion of extant serotypes uncommon prior to vaccination rather than capsule-switching drove the increased frequency of NVT serotypes post-vaccination. The majority of the capsule-switches occurred pre-vaccination but the specific times when those events occurred could not be established due to short sampling frame, which could bias temporal phylogenetic signal [2 years] (Fig 3, S8 Fig, S3 Table). Six capsule-switch events were detected based on phylogenetic clustering and ST profiles namely serotype 11A to 20 in GC7, 13 to 19A in GC9, 16F to 19F in GC13, 9V to 28F in GC21, and 7C to NT in GC23 (S8 Fig). The capsule-switched strains circulated at low frequency (<1%) both pre‐ and post-vaccination ruling them out as a driver for the NVT serotype replacement. Of these capsule-switched serotypes, only serotype 28F, which was not detected prior to vaccination in carriage and previously studied invasive datasets [19], underwent a significant clonal expansion post-vaccination (Fig 2e,f and Fig 3). Of the two serotype 28F lineages, the increase of serotype 28F strains was due to clonal expansion of strains in NVT GC2 rather than the serotype 9V to 28F vaccine-escape capsule-switched variants in GC21 (Fig 2e,f). No serotype 28F strains were detected pre-vaccination including in previous IPD datasets, which suggested either circulation at undetectable levels prevaccination or importation from other countries.

**Fig 3:**
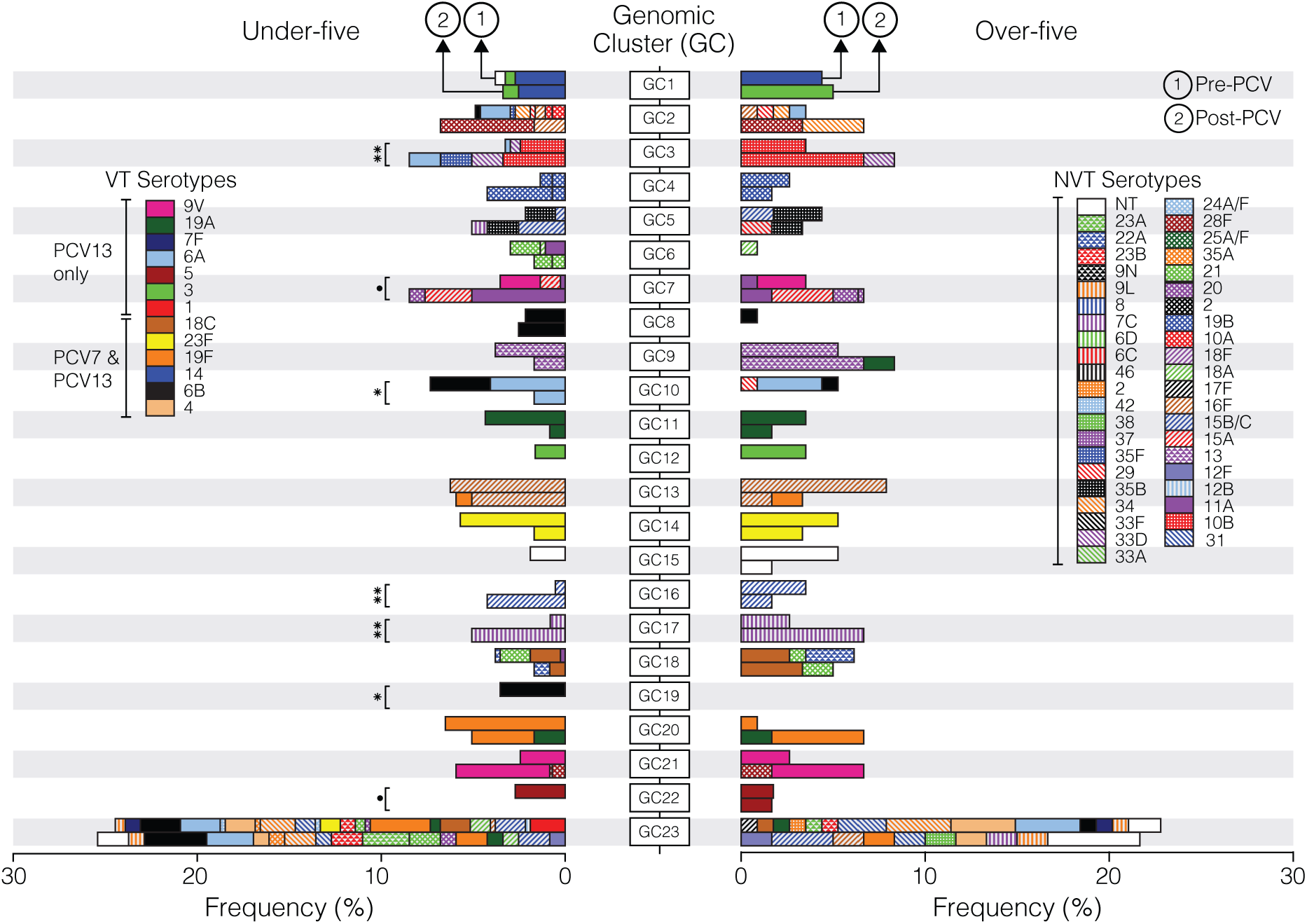
Dynamics of pneumococcal GCs and serotypes. The leftward facing stacked bar graph shows frequency of GCs in under-fives while (b) the rightward facing bar graph shows frequency of GCs and their constituent serotypes in over-fives before and after PCV introduction. The bar graphs are aligned by genomic clusters (GC) for easy comparisons of frequency of serotypes pre‐ and post-vaccination between the two age groups. The serotypes are distinguished by different colours in the bar graphs as described in the key. GC23 is the ‘bin’ cluster consisting of unclustered strains not placed in clusters GC1-22. The GCs whose frequency changed significantly post-vaccination are marked with asterisks: *P*<0.05 (*) and *P*<0.01 (**) and those with borderline significance *P*<0.095 (.). The Fisher’s exact test was used to determine *P*-values.

**Fig 4:**
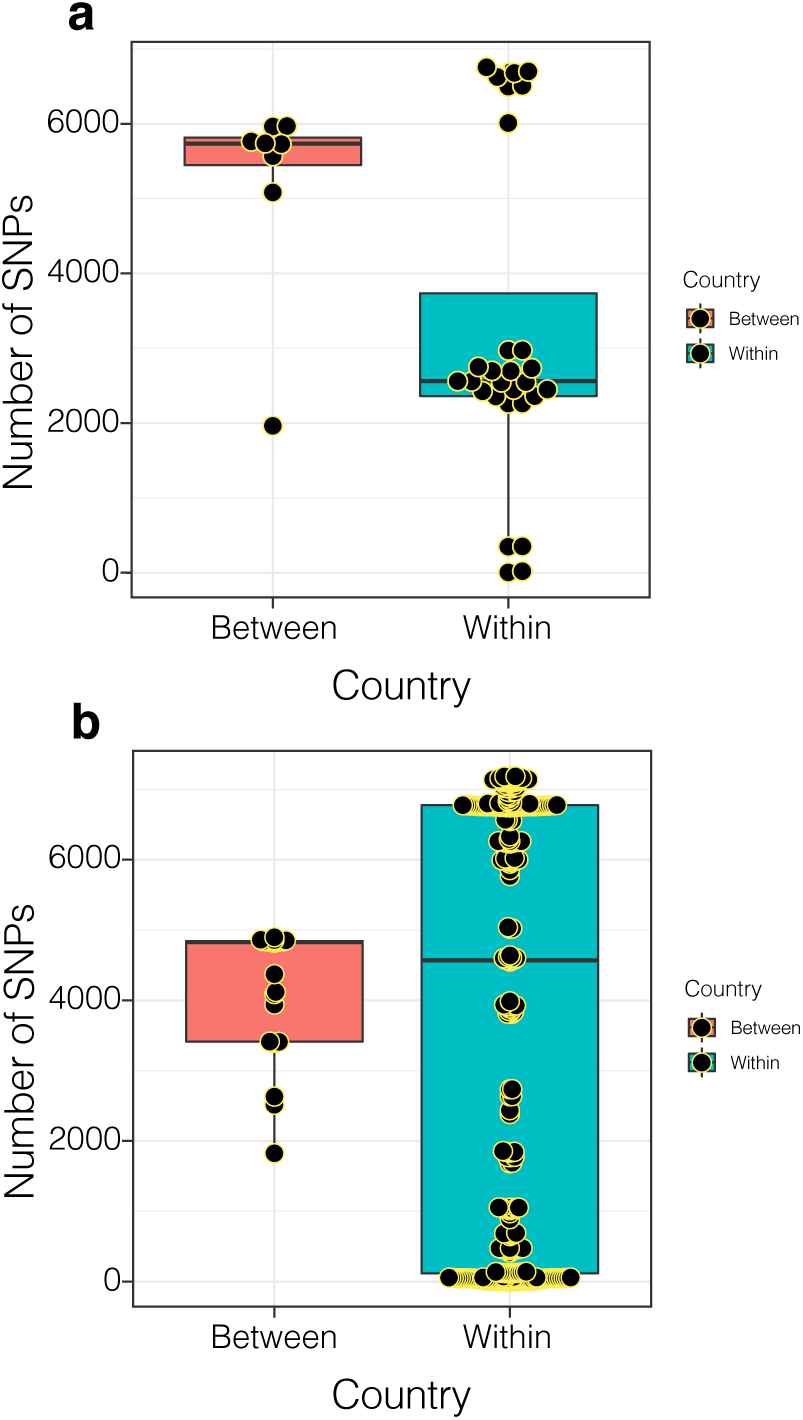
Genetic diversity of a recently emerged serotypes (28F). Boxplots showing within (Malawi) and between country (Malawi and South Africa) genetic diversity of serotype 28F strains showing in (a) GC2 (b) GC21. Lineage GC21 also include serotype 9V strains, which some of which underwent a capsule-switch to acquire a serotype 28F capsule. Additional details are provided in in S11 Fig

Additional strains with similar ST profiles were searched in the global collection of ≈20,000 strains with similar ST profiles to investigate whether there was importation of serotype 28F strains from other countries. Only two strains were identified from South Africa, a serotype 28F and 9V clustering with strains in GC2 and GC21 respectively both identified only post-13 introduction in South Africa (2013 and 2014). The closest matching strains from our setting were distinguished from the South African strains by 1,966 and 2,284 SNPs in GC2 and GC21, which implied no recent importation post-vaccination (Fig 4a,b). Furthermore, the maximum sequence divergence between serotype 28F strains in GC2 was 6,757 SNPs, which is a further contradiction of the hypothesis that serotype 28F was imported post-vaccination because recently imported clones typically exhibit less diversity caused by transmission bottleneck. With such a short time frame from PCV introduction (2 years), this would have been insufficient for the imported strains to accrue such high genetic diversity considering that the pneumococcal mutation rate is ≈4 SNPs per year [20]. Furthermore, although the capsule-switched 28F strains in GC21 clustered together with the serotype 9V strains, these serotypes had different STs, which suggests that the capsule-switch event occurred earlier prior to vaccination. Therefore, these findings demonstrate that newly emerging serotypes post-vaccination were not due to importation of novel serotypes from other countries but rather expansion of extant serotypes circulating at undetectable levels prior to vaccination.

### Reduction of Simpson diversity index indicates PCV impact

Reduction in Simpson diversity among VT serotypes post-vaccination occurred in under-fives (*P*=0.022, resampling) further supporting direct PCV effect (Fig 5a, S6 Table). Reduction in Simpson diversity was also detected in NVT strains from under-fives but to a lesser extent while the opposite trend occurred in NVT strains from over-fives (S10a,b Fig, S7 Table), implying incomplete serotype replacement by NVT serotypes post-vaccination. Pre-vaccination diversity was similar for VT and NVT strains but following vaccination Simpson diversity was higher in NVT than VT strains following vaccination (*P*=0.004, resampling) (Fig 5b). The Simpson diversity index was higher for STs than serotypes both before (*P*=0.011, resampling) and after vaccination (*P*=0, resampling) (S9 Fig). No changes in Simpson diversity were detected for the composition of STs post-vaccination (S10 Fig, S6,7 Table). The high stability of Simpson diversity in NVT strains post-vaccination could imply limited serotype replacement by NVTs, which has resulted in incomplete restoration of the serotype diversity to the levels observed pre-vaccination following vaccine-induced clearance of VT strains.

**Fig 5:**
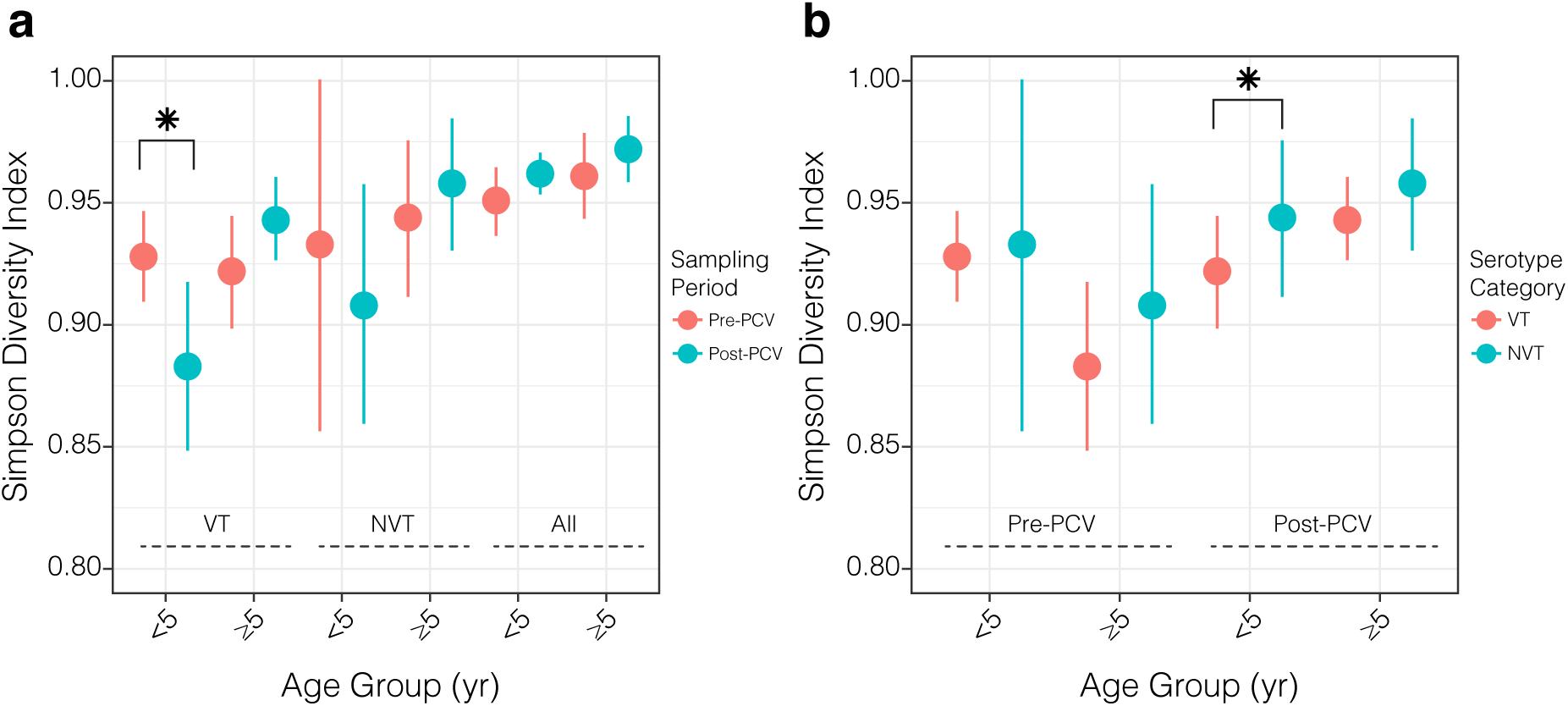
Serotype composition and diversity in context of PCV. (a) Simpson diversity index for composition of serotypes between pre‐ and post-vaccination datasets among VT, NVT and all strains among isolates. (b) Simpson diversity index for composition of serotypes between VT and NVT strains sampled pre‐ and postvaccination. Statistically significant changes are marked with asterisks: ‘ns’: not significant, *P*<0.05 (*), *P*<0.01 (**) and *P*<0.001 (***). The estimates and *P*-values for frequency of VTs and Simpson diversity are summarised in S6,7 Table.

### Antibiotic resistance

An important set of pneumococcal accessory genes include those encoding proteins that confer resistance to antibiotics. Previous work [21] showed almost universal resistance and complete sensitivity of pneumococcal strains to co-trimoxazole and ceftriaxone respectively therefore we did not investigate these antibiotics. Resistance rates were detected genotypically by quantifying the frequency of the antibiotic resistance conferring genes namely chloramphenicol acetyltransferase encoding gene (*cat*_pC194_) for chloramphenicol, macrolide efflux pump encoding genes (*mefA* and *mefE*) and ribosomal RNA methyltransferase (*ermB*) for erythromycin, ribosomal protection protein-encoding gene (*tetM*) for tetracycline. For penicillin, the minimum inhibitory concentrations (MIC) were genotypically predicted using allelic variation in the transpeptidase domain of the penicillin binding proteins (PBP) from which the binary resistant-susceptible phenotype was inferred using BSAC criteria [22] (Fig 6a-d). There were no significant changes in resistance rates against the four antibiotics (Fig 6e-h). Interestingly, despite no change in penicillin resistance rate post-vaccination, the MICs decreased significantly (*P*=0.0098, Student’s t test) postvaccination due to vaccine-induced clearance of intermediate resistant lineages particularly serotype 3 in GC12 (Fig 6i, S8 Table). This exemplifies how vaccine usage can be strategically employed to clear not only highly prevalent and antibiotic resistant pneumococcal lineages globally but also intermediate resistant lineages with high likelihood to express full resistance before they do [23]. Further genomic analysis revealed the existence of a diverse catalogue of mobile genetic elements (MGE) which disseminated genes responsible for resistance against macrolides, tetracycline and chloramphenicol antibiotics (Fig 6j). The macrolide (erythromycin) resistance conferring genes were associated with Tn*916*, Tn*6003*, Tn*2009* and Tn*2010* elements but not Tn*1545*, which is common elsewhere [24] (Fig 6c,d). The *mefA/E* gene were disseminated by Tn*2009* and Tn*2010*-like elements while ermB was carried by Tn *6003* and Tn916-like conjugative elements (Fig 6c,d). Both cat_pC194_ and *tetM* genes were located on Tn*5253*-like conjugative elements but tetM was more common due to its association with additional independent elements mainly Tn*5251* and Tn*2009*-like elements (Fig 6c,d). These findings suggests no significant post-vaccination perturbation of the accessory gene pool associated with antibiotic resistance.

**Fig 6:**
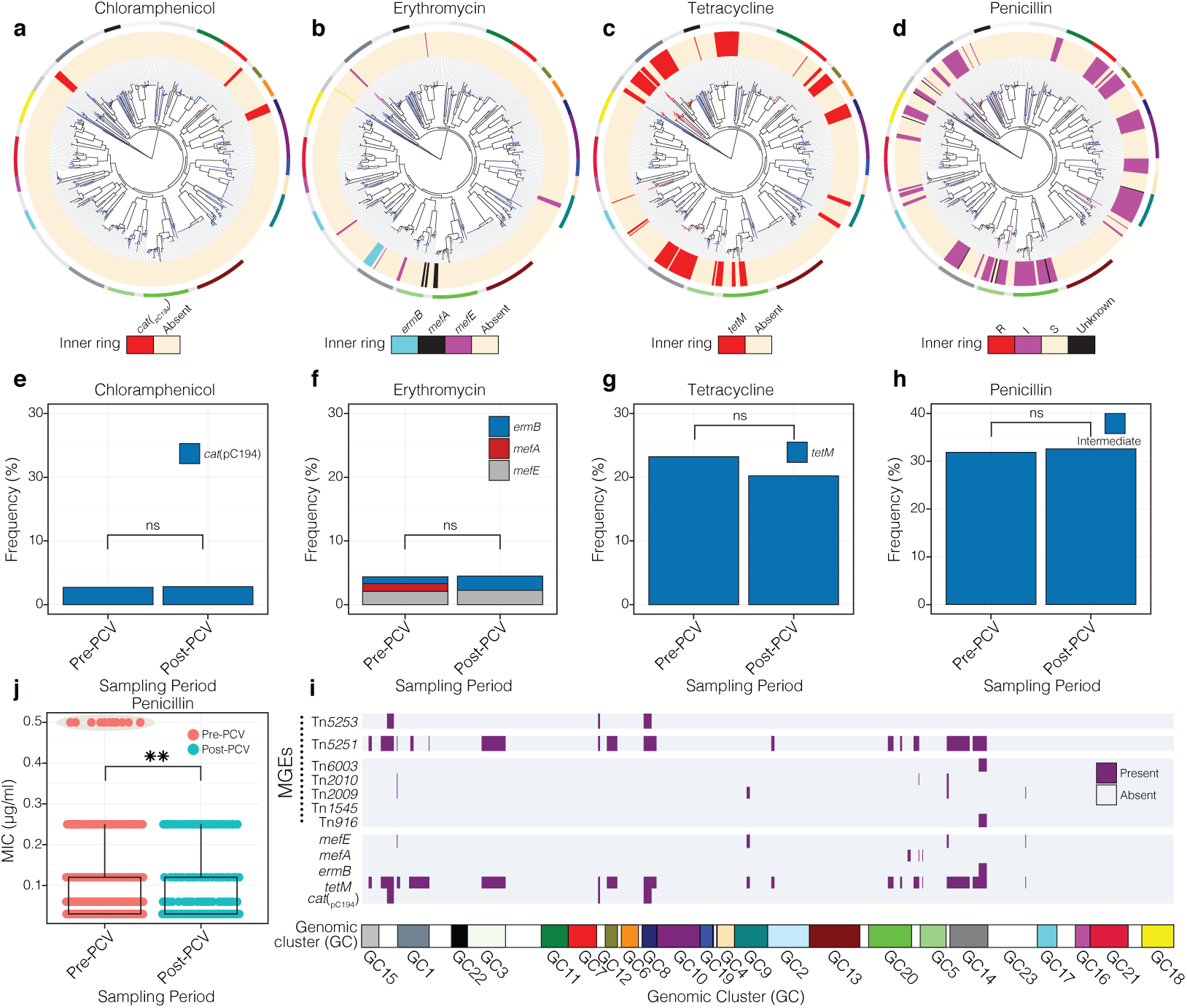
Distribution of antibiotic resistance genes and MGEs. (a) Distribution of *cat*_pC194_ chloramphenicol resistance gene. (b) Distribution of *mefA*, *mefE* and *ermB* erythromycin resistance genes. (c) Distribution of *tetM* tetracycline resistance gene. (d) Distribution of penicillin MICs. Branches of the maximum likelihood phylogenies and innermost ring surrounding the trees in a-c are coloured by presence and absence of the genes as shown in the key at the bottom of the phylogeny. The outermost ring around the phylogenies shows the GCs corresponding to those in Fig 1b and the colour strip at the bottom of this figure. (e-h) Frequency of genotypic antibiotic resistance rates for chloramphenicol, erythromycin, tetracycline and intermediate penicillin resistance pre‐ and post-vaccination. (i) Distribution of penicillin MICs pre‐ and post-vaccination. (j) The horizontal panel shows the distribution of the antibiotic resistance conferring genes and MGEs that disseminate them. The subsets with statistically significant changes are marked with asterisks: ‘ns’: not significant, *P*<0.05 (*), *P*<0.01 (**) and *P*<0.001 (***).

### Vaccine-induced accessory genome dynamics

Distribution of intermediate frequency accessory genes detected in 5% to 95% of the strains were similar between different subsets of isolates including those from over-fives and under-fives regardless of serotype category both pre‐ and post-vaccination (Fig 7a, S12 Fig). No significant differences in frequency of accessory genes were detected among VT strains pre‐ and post-vaccination while three accessory genes showed significant change among NVT strains following vaccination (Fig 7a,b). The accessory genes which changed in frequency post-vaccination were detected by fitting a logistic regression model for the binary presence-absence genotype for each gene and assessing the coefficients for the sampling period. Other variables namely age group and serotype category were included in the model formulation to account for their group effects. Similar analysis was done to determine accessory genes associated with different age groups and serotype categories. Bonferroni adjustment was done to control for multiple testing and no genes were associated with specific age group (Fig 7d) but as expected many genes (≈400) were associated with different serotype categories (Fig 7e). Forty-two accessory genes showed significant change in frequency post-vaccination, of which approximately half (52.38%) increased postvaccination (Table 1, Fig 7f, S9 Table). The accessory genes that increased postvaccination encoded for a glycosyl transferase (odds ratio[OR]=3.34, *P*=9.71×10^−6^), bacteriolysin family protein (OR=3.34, *P*=3.46×10^−5^), type I restriction modification system (RMS) R subunit (OR=3.34, *P*=3.46×10^−5^) and other diverse functions including sugar transport, MGE and phage activity. Conversely, genes whose frequency decreased were associated with bacteriocin gene *blpQ* (OR=0.19, *P*=5.20×10^−6^) and other genes associated with bacteriocins, conjugative elements, two-component systems and phages (all OR≈0.41, *P*<1.52×10^−2^). Interestingly, a capsule biosyn-thesis gene (*wzx*) encoding a repeating unit of a flippase protein also showed a decreasing frequency (0R=0.43, *P*=0.042). Other genes with significant changes in frequency encoded proteins with a diverse functional repertoire but notably included MGEs [phages/transposase/insertion elements] (11/42), bacteriocins (3/42), RMS systems (2/42) and competence (2/42), which have been recently implicated to play a role in negative frequency dependent selection (NFDS) [25]. These findings indicate that PCV has perturbed pneumococcal population not only at serotype level but also accessory genome level.

**Fig 7:**
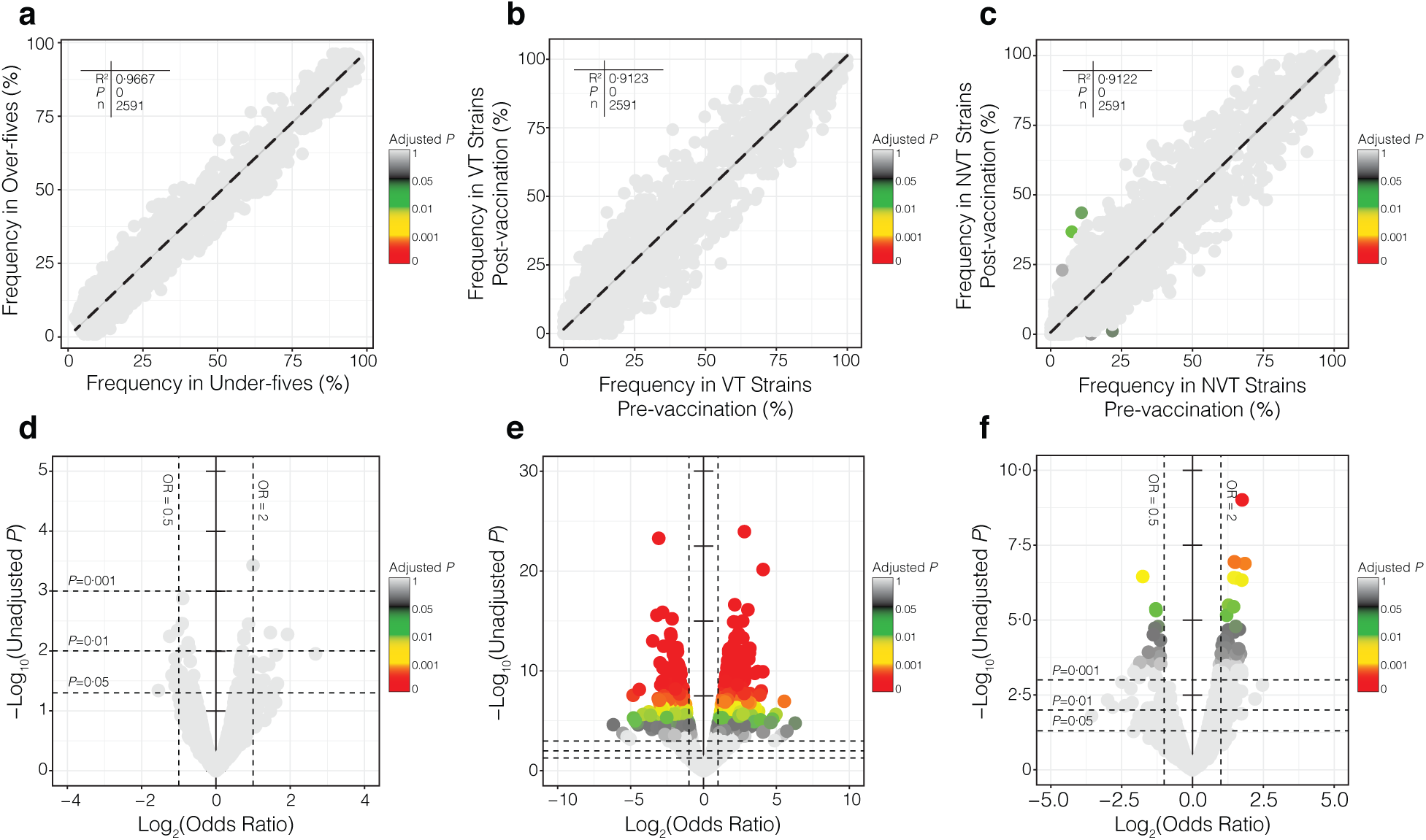
Pneumococcal accessory genome dynamics. The distribution of 2,591 intermediate frequency accessory genes in the entire pneumococcal population. (a) Scatter plot showing frequency of accessory genes between isolates sampled from under-fives and over-fives pre-vaccination. (b) Scatter plot showing frequency of genes among VT isolates pre‐ and post-vaccination. (c) Scatter plot showing frequency of genes among NVT isolates pre‐ and post-vaccination. Coefficients from linear regression and are labelled on the plots. Volcano plots show magnitude (log_2_ odds ratio) on the x-axis and statistical significance (-log_10_ *P*-value) for *P*-values and odds ratio for the association of accessory genes with different variables namely (d) age group, (e) serotype category and (f) sampling period. The points were coloured by adjusted *P*-values after correcting for multiple testing using Bonferroni method.

**Table 1:** Coefficients for the effect of sampling period on presence and absence of accessory genes using logistic regression accounting for age and serotype category after Bonferroni adjustment for multiple testing (only top 25 hits shown). Full list is shown in S9 Table.

| Accessory gene | Odds ratio (95% CI) | Raw *P*-value | Adjusted *P*-value | Description/product |
| --- | --- | --- | --- | --- |
| COG_6818 | 3.34 (2.24,4.99) | 3.75×10<sup>-09</sup> | 9.71×10<sup>-06</sup> | Glycosyl transferase |
| COG_7104 | 3.34 (2.20,5.06) | 1.33×10<sup>-08</sup> | 3.46×10<sup>-05</sup> | Double glycine cleavage site bacteriolysin superfamily |
| *hsdR_1* | 3.34 (2.20,5.06) | 1.33×10<sup>-08</sup> | 3.46×10<sup>-05</sup> | Type I restriction-modification system, R subunit |
| COG_2038 | 2.78 (1.91,4.06) | 1.12×10<sup>-07</sup> | 2.91×10<sup>-04</sup> | ABC transporter permease |
| COG_900 | 2.78 (1.91,4.06) | 1.12×10<sup>-07</sup> | 2.91×10<sup>-04</sup> | IS630-Spn1, transposase *Orf1* |
| COG_5542 | 2.84 (1.93,4.18) | 1.17×10<sup>-07</sup> | 3.03×10<sup>-04</sup> | Replication protein |
| *ptrB* | 0.30 (0.19,0.47) | 3.48×10<sup>-07</sup> | 9.02×10<sup>-04</sup> | Prolyl oligopeptidase family protein |
| COG_4000 | 2.95 (1.94,4.48) | 3.96×10<sup>-07</sup> | 1.03×10<sup>-03</sup> | Phage protein gp27 |
| *rlmN* | 2.95 (1.94,4.48) | 3.96×10<sup>-07</sup> | 1.03×10<sup>-03</sup> | Ribosomal RNA large subunit methyltransferase N |
| COG_4863 | 3.34 (2.09,5.33) | 4.59×10<sup>-07</sup> | 1.19×10<sup>-03</sup> | Hypothetical protein |
| COG_5488 | 2.54 (1.76,3.65) | 5.24×10<sup>-07</sup> | 1.36×10<sup>-03</sup> | Phage protein |
| COG_842 | 0.40 (0.28,0.58) | 6.98×10<sup>-07</sup> | 1.81×10<sup>-03</sup> | ABC transporter permease |
| IS861 truncation | 0.40 (0.28,0.58) | 6.98×10<sup>-07</sup> | 1.81×10<sup>-03</sup> | IS861 truncation |
| COG_1881 | 2.74 (1.82,4.14) | 1.59×10<sup>-06</sup> | 4.11×10<sup>-03</sup> | SNF2 family protein |
| COG_802 | 2.74 (1.82,4.14) | 1.59×10<sup>-06</sup> | 4.11×10<sup>-03</sup> | Transposase |
| COG_4974 | 3.00 (1.91,4.72) | 1.93×10<sup>-06</sup> | 5.00×10<sup>-03</sup> | Membrane associated protein |
| COG_1144 | 2.41 (1.67,3.50) | 3.15×10<sup>-06</sup> | 8.17×10<sup>-03</sup> | 2-isopropylmalate synthase |
| COG_3978 | 0.41 (0.28,0.60) | 4.17×10<sup>-06</sup> | 1.08×10<sup>-02</sup> | Rep protein |
| COG_5509 | 0.41 (0.28,0.60) | 4.17×10<sup>-06</sup> | 1.08×10<sup>-02</sup> | Phage protein gp27 |
| COG_6116 | 0.41 (0.28,0.60) | 4.17×10<sup>-06</sup> | 1.08×10<sup>-02</sup> | ABC transporter permease |
| *saeS* | 0.41 (0.28,0.60) | 4.21×10<sup>-06</sup> | 1.09×10<sup>-02</sup> | Histidine kinase |
| COG_6305 | 0.41 (0.28,0.60) | 4.77×10<sup>-06</sup> | 1.24×10<sup>-02</sup> | Bacteriocin |
| COG_748 | 0.41 (0.28,0.60) | 4.77×10<sup>-06</sup> | 1.24×10<sup>-02</sup> | Hypothetical protein |
| *blpPQ* | 0.19 (0.09,0.39) | 5.20×10<sup>-06</sup> | 1.35×10<sup>-02</sup> | Bacteriocin BlpPQ |
| COG_2726 | 0.41 (0.28,0.61) | 5.84×10<sup>-06</sup> | 1.51×10<sup>-02</sup> | Tn5252, *Orf 9* protein |

## Discussion

The evolution of the pneumococcus in carriage is ever-changing therefore sampling this niche provides unrivalled opportunities to monitor and keep track of ongoing genomic adaptations over-time [5, 7]. Our genomic study demonstrates this by demonstrating changes in pneumococcal population structure following the introduction of PCV vaccine in previously vaccine-naïve low-income setting where, unlike in higher income settings such information is usually unavailable despite higher disease burden [26]. In contrast from other studies, our dataset is unique because it includes strains from both children and adults, which provides a more an opportunity to investigate PCV-induced population structuring in both vaccine-eligible and ineligible individuals. Our findings reveal changes at serotype, lineage and accessory genome level caused by vaccine-induced population restructuring specifically due to substantial clearance of VT serotypes in vaccinated under-five population but not unvaccinated over-five population. This reflects a direct consequence of PCV but limited and possibly delayed indirect protection in contrast to other settings e.g. UK where population herd immunity is higher albeit epidemiological differences with our population [3, 10]. Further ongoing and future studies will investigate factors mitigating herd effects, which may include higher HIV burden and vaccine scheduling. Lineage-specific changes revealed subtle serotype dynamics associated with clearance, clonal expansion and emergence of serotypes. Furthermore, while there was no change in the accessory gene pool associated with antibiotic resistance, there significant changes in other genes typically those encoding MGEs and bacteriocins. Additional statistical modelling shows that other accessory genes have undergone significant changes in frequency due to the population perturbations, which could be temporary and may return to equilibrium gene frequency due to NFDS [25]. Altogether, these findings reveal an early impact of PCV on serotype and lineage distribution, and accessory gene dynamics in pneumococcal carriage post-vaccination.

The impact of PCV is shown by the reduction in frequency of VT serotypes, which occurred only in the largely vaccinated under-five population but not older and unvaccinated individuals. This implied high direct impact but limited indirect protection via herd immunity. While the frequency of VTs was higher in under-fives than over-fives, following vaccination these converged to the same amount (33%) but whether this is coincidental or represents some unknown phenomena threshold for VT frequency in the population remains unknown. Further assessment of Simpson diversity in serotype composition in VT and NVT strains provided further insights on the strain dynamics. Firstly, the Simpson diversity for ST composition remained largely unchanged post-vaccination [27, 28]. This suggests PCV did not significantly disrupt the ST composition in a similar fashion to the way it did for serotype composition possibly because of higher (≈3 times) saturation of STs than serotypes in pneumococcal population. Secondly, there was an increase of the diversity in NVT strains after vaccination albeit not statistically significant but serotype diversity was not restored to the levels observed pre-vaccination before clearance of VT serotypes, which implied that serotype replacement was incomplete. How serotype replacement in our setting will compare with other settings where settings remains to be fully understood once the population has reached a new equilibrium and replacement is complete.

Consistent with data from other settings [13], the observed modest serotype replacement has been driven by clonal expansion of extant serotypes prior to vaccination, which were masked due to competition by their VT counterparts. Unlike post-PCV7 introduction where serotype 19A was the dominant replacement serotype, in our setting replacement is driven by multiple serotypes including serotype 7C, 23A, 15B/C and 28F rather than a single dominant NVT clone. Furthermore, while certain capsule-switched strains were detected post-vaccination but whether these occurred postvaccination due to vaccine pressure is unknown but there was evidence that the majority of these were likely extant prior to vaccination but at undetectable frequencies such as the serotype 9V to 28F vaccine-escape capsule-switched strains. Emergence of novel serotypes post-vaccination was uncommon with only serotype 28F detected post-vaccination only but this clone showed high sequence diversity and was not genetically similar to strains from other countries suggesting not only no recent importation post-vaccination but also circulation at undetectable levels prior to vaccination. Further studies are needed to assess the degree with which such replacement serotypes will cause disease in our setting and globally [29]. Taken together, these findings suggest there is incomplete serotype replacement in our setting after two years from PCV introduction mostly in under-fives compared to older age groups.

Clinically, the stability of the antibiotic resistance rates is not concerning considering that resistance rates were already lower than in IPD [19, 21, 30]. However, consistent with findings elsewhere [31, 32], the significant decrease of intermediate-penicillin resistance rate exemplifies the advantages of PCVs when strategically harnessed to thwart further emergence and expansion of clones with intermediate resistance to antimicrobials before they higher resistance is achieved [31]. While the accessory gene pool associated with antibiotic resistance did not change significantly, the changes detected in other accessory genes using logistic regression signaled that the pneumococcal population has been perturbed and these could likely indicate genes with the potential to drive NFDS. The perturbed-frequency genes revealed diverse functions with the majority associated with MGEs possibly reflecting their rapid mobility between pneumococcal strains. Other genes were associated with capsule biosynthesis (*wzx*) while those encoding for bacteriocin activity associated proteins important in mediating strain competition dynamics were also possibly under selection [33, 34]. These finding provide a clear evidence that PCV has induced population changes not only at serotype but also accessory genome level but these changes did not seemingly favour higher antibiotic resistance. The resultant phenotypic changes associated with the perturbed accessory genes and allelic variation in the core gene previously shown to drive metabolic shifts in strains post-vaccination warrants further investigation [35].

We have provided clear evidence of vaccine-induced perturbed pneumococcal population at serotype, lineage and accessory genome level after only two years following the introduction of PCV in a pneumococcal vaccine-naïve setting in Northern Malawi. This reflects a PCV impact, which is largely restricted to vaccinated under-fives but not unvaccinated older individuals highlighting limited manifestation of population herd immunity. Our data is timely and improves our understanding of the impact of PCV on pneumococcal population structure in high-disease burden but low-income settings. Our data revealed a perturbed pneumococcal carriage population after two years from PCV introduction but continued assiduous surveillance and WGS remain crucial to adequately monitor long-term effects of PCV particularly after the equilibrium population dynamics have been re-established. Together with findings gained from surveillance and clinical trials of PCVs across SSA and globally, our findings will inform future conjugate vaccine design strategies and how their beneficial effects can be maximised especially in the most vulnerable tropical populations.

## Materials and methods

### Study population and isolate selection

A subset of 660 pneumococcal isolates collected through multiple household surveys were selected for WGS (S1 Data). These isolates were obtained from nasopharyngeal carriage in healthy children and adults in Karonga district of northern Malawi before vaccination in 2009 (n=370) and 2010 (n=112), and two years after vaccination in 2014 (n=178) were selected for WGS. The strains were representative of nasopharyngeal swabs and were samples from individuals from different age groups. By age group, 376 strains were sampled from the under-fives (<5 years old) and 130 strains from over-fives (≥5 years old) before vaccination while 106 and 48 strains were obtained from under-fives and over-fives post-vaccination. The nasal swabs were stored and processed as previously described [36]. The ethical approvals for the study were granted by National Health Sciences Research Committee in Malawi (NHSRC 490) and University of Malawi College of Medicine Research Ethics Committee (P.O8/14/1614).

### Genomic DNA preparation and sequencing

Genomic DNA extraction was done using QIAamp DNA mini kit (Qiagen, Hilden, Germany), QIAgen Biorobot (Qiagen, Hilden, Germany) and Wizard^®^ DNA Genomic DNA Purification Kit (Promega, Wisconsin, USA) as previously described [37]. Preparation of genomic DNA libraries and sequencing was done at Wellcome Sanger Institute using Illumina Genome Analyzer II and HiSeq platforms (Illumina, CA, USA). The length of reads ranged between 100 and 125 bases (median: 100) with a mean quality of 35.32 (S1 Fig) while mean number of mapped reads was 4,068,781 reads per isolate (S2 Fig). De novo sequence assemblies were generated using a pipeline [38], which uses Velvet v1.2.09 [39] and VelvetOptimiser v2.2.5 [40] (k-mers between 66.0% to 90.0% of the read length) to assemble reads, SSPACE Basic v2.0 [41] for assembly scaffolding (≈16 iterations), GapFiller v1.10 [42] for assembly gap closing and SMALT v0.7.4 (www.sourceforge.net/projects/smalt/) to re-map reads to the assembly. The mean genome size, contig length and number of contigs were 2,125,143bp, 54,071bp and 48 contigs respectively while the mean N50 values was 129,431bp (S3,4 Fig). The sequence reads were deposited in the European Nucleotide Archive (S1 Data).

### Capsule and sequence typing

The capsule types (serotypes) were defined using an *in silico* typing approach [43], which maps short sequence reads against reference sequences of the capsule-encoding polysaccharide synthesis (CPS) loci, which determines serotypes based antibody binding to its antigens [44]. The sequence types (ST) were inferred from assemblies based on loci of seven housekeeping genes for pneumococcal multilocus sequence typing (MLST) scheme [45] using MLSTcheck [46].

### Presence of antibiotic resistance genes

The presence of antibiotic resistance genes for different antibiotics (tetracycline, chloramphenicol and erythromycin) and mobile genetic elements that disseminate them were investigated using nucleotide-BLAST v2.2.30 [47] with E-value of <0.001, sequence coverage >80% and nucleotide identity >80%. For penicillin whose resistance is caused by chromosomal mutations unlike genes disseminated by mobile genetic elements (MGE), the minimum inhibitory concentrations (MIC) were genotypically predicted using a robust analysis pipeline developed by the Centers for Disease Control and Prevention (CDC), which uses allelic variation in the transpeptidase domain of the penicillin binding proteins (PBP) - PBP1a, PBP2b and PBP2x to infer MICs [22]. The inferred MICs were translated into binary resistant-susceptible phenotype using the British Society for Antimicrobial Chemotherapy (BSAC) breakpoint [48].

### Gene annotation, alignment and population structure

The sequenced and assembled genomes were annotated using Prokka v1.11 [49]. We identified core and accessory genes, present in ≥99% and <99% of the strains respectively, by clustering coding sequences using Roary v3.6.1 pan-genome pipeline [50]. The core‐ and pan-genome were comprised of 660 and 9,472 genes respectively and a 1,050,021bp core-genome alignment with 88,961bp single nucleotide polymorphism (SNP) positions identified using Snp-Sites v2.3.2 [51] was generated from Roary analysis (S5 Fig). The core SNPs were clustered into genomic clusters (GCs) using the hierarchical clustering module (hierBAPS) in BAPS v6.0 [18]. DNA sequence translation and format conversion was done using BioPython [52] while nucleotide-BLAST v2.2.30 [47] and ACT v13.0.0 [53] was used for sequence comparison.

### Phylogenetic tree construction

Maximum likelihood phylogenetic trees of the strains was constructed from the core gene alignment using FastTree-SSE3 v2.1.3 [54]. The GC or lineage-specific trees were constructed using RAxML v7.0.4 [55] from whole genome alignments after removing regions with putative recombination events using Gubbins v1.4.10 [56]. The phylogenetic trees were generated using a general time reversible (GTR) model [57], Gamma heterogeneity between nucleotide sites [58] and 100 bootstrap replicates parameters [59]. The phylogenetic tree was rooted at the midpoint of the branch separating most divergent strains. Visualisation of the tree Together with the strain’s metadata was done using iToL v2.1 [60].

### Statistical analysis

The changes in frequency of serotypes, antibiotic resistance and accessory genome content were assessed using Fisher’s Exact test. Odds ratios for detecting GCs and serotypes pre‐ and post-vaccination were determined (pseudo-counts of 1 to avoid division by zeros). Changes in the composition of serotypes and STs were detected by using Simpson diversity index and the *P*-values were detected by resampling using Jackknife approach (www.comparingpartitions.info). Logistic regression was used to assess changes in gene presence-absence patterns of the intermediate frequency accessory genes (present at frequency from 5% to 95%) before and after vaccination while controlling for the effects of age group and serotype category. The reference levels for each variable in the regression were as follows: ‘pre-vaccination’ for sampling period, ‘over-five’ for age group and ‘NVT’ for serotype category. The estimated coefficients for each variable were extracted and summarised and *P*-values were for each gene were adjusted using Bonferroni correction to account for multiple comparisons. The statistical analysis was done using R v3.1.2 (R Core Team, 2013), GraphPad Prism v7.0 (GraphPad Software, California, USA) and Python v2.7.9 (Python Software Foundation).

### Ethics statement

Written informed consent was obtained from adults while parents, guardians and caregivers of child participants. The ethical approvals for the study were granted by National Health Sciences Research Committee in Malawi (approval #: NHSRC 490 and 1232), the London School of Hygiene and Tropical Medicine (approval #5345) and the University of Liverpool (approval #: 670) and University of Malawi College of Medicine Research Ethics Committee (approval #: P.O8/14/1614). Nasopharyngeal swabs were collected from healthy children and adults as previously described and the samples were de-identified in the analysis.

## Author contributions

C.C., N.F., D.B.E. and S.D.B. conceived and designed the study. N.F., D.B.E. and S.D.B. supervised the study. N.F., D.B.E., L.M., R.F.B. and S.D.B. secured funding. E.H., T.T. and N.F. collected samples. M.A., A.W.K., J.E.C. and C.P. performed molecular and microbiology experiments. S.D.B. supervised whole genome sequencing and genomic analysis. C.C. and R.A.G. checked quality of the sequence assemblies. C.C. performed genomic and statistical analyses. P.M. assisted with data analysis and interpretation. C.C. and S.D.B. wrote initial draft of the paper. L.M., R.F.B., A.K., W.P.H. and R.S.H. contributed to data interpretation. All authors contributed to writing and reviewing of the paper.

## Acknowledgements

We acknowledge work by clinical and laboratory staff at Malawi Epidemiology and Intervention Research Unit (MEIRU) and Malawi-Liverpool-Wellcome Trust Clinical Research Programme (MLW) who collected and prepared the samples respectively, and sequencing and informatics teams at Wellcome Sanger Institute. This work was funded by Bill and Melinda Gates Foundation (grant #OPP1034556) and Wellcome UK (grant #084679/Z/08/Z). C.C. acknowledge PhD studentship funding from Commonwealth Scholarship Commission. The contents of this paper are solely responsibility of the authors and does not necessarily represent official views of their affiliated institutions and funding agencies.

